# Implications of Spike Protein Interactions with Zn-bound form of ACE2: A Computational Structural Study

**DOI:** 10.1101/2022.06.05.494906

**Authors:** Peter R. Fatouros, Urmi Roy, Shantanu Sur

**Affiliations:** Department of Chemical and Biomolecular Engineering, Clarkson University, 8 Clarkson Avenue, Potsdam NY 13699, United States; Department of Chemistry and Biomolecular Science, Clarkson University, 8 Clarkson Avenue, Potsdam NY 13699, United States; Department of Biology, Clarkson University, 8 Clarkson Avenue, Potsdam NY 13699, United States

**Keywords:** binding free energy, SARS-CoV-2, spike protein, structural dynamics, zinc-bound ACE2, zinc parameterization

## Abstract

The COVID-19 pandemic has generated a major interest in designing inhibitors to prevent SARS-CoV-2 binding on host cells to protect against infection. One promising approach to such research utilizes molecular dynamics (MD) to identify potential inhibitors that can prevent the interaction between spike (S) protein on the virus and angiotensin converting enzyme 2 (ACE2) receptor on the host cells. In these studies, many groups have chosen to exclude a zinc (Zn) ion bound to the ACE2 molecule which is critical for enzymatic activity. While the relatively distant location of Zn ion from the S protein binding site (S1 domain), combined with the difficulties in modeling this ion have motivated the decision of exclusion, Zn can potentially contribute to the structural stability of the entire protein, and thus, may have implications on spike protein interaction. In this study, we explored the effects of excluding Zn on the structural stability and binding free energy of the ACE2-S1 protein complex. We generated two versions of an experimentally-derived structure of the ACE2-S1 protein complex: one with Zn and one without. Examining the differences between these two complexes during MD simulation, we found that the Zn-bound complex exhibited greater instability at nearly all residues except for the interacting residues, which were more stable in the Zn-bound complex. Additionally, the Zn-bound complex had a stronger binding free energy at all internal dielectric constants greater than one. Since binding free energy is often used to score inhibitors’ performances, excluding Zn could potentially have implications on inhibitor selection and performance, both in the ACE2-S1 protein system and other protein complexes that include the Zn ion.

## Introduction

The global impact of the COVID-19 pandemic has led to extensive research on prevention of infection through development of effective vaccines and inhibitors. While some groups have taken an experimental approach to search for novel inhibitory molecules, many others have used computational methods due to their time and cost effectiveness (1). The research on inhibitory molecules for SARS-CoV-2 has been directed to two main targets, the S1 domain of spike (S) protein on the virus and the angiotensin converting enzyme 2 (ACE2) protein on the host cells, which serves as a binding site for S protein (2) (3). In the computational studies, a wide array of potential inhibitors are docked to either S1 or ACE2 and are then evaluated based on the thermodynamics and/or the kinetics of their interaction. While computational methods can allow researchers to evaluate many potential inhibitors in a relatively short period of time, selection of appropriate computational models are critical to generate results that closely reflect the experimental data.

Many receptors and enzymes contain metals ions such as zinc and copper, serving an important role in the structure and/or function of these protein molecules. The abundance of metalloproteins is extremely common and it is estimated that nearly half of all human proteins contain at least one metal ion (4). These metal ions are often essential for catalytic or co-catalytic activity, and can have large implications on the overall structural stability of the protein (5). Various functional groups within the protein can coordinate to metal ions, which form strong bonds that increase the structural stability within the region. Additionally, metal ions can have far-reaching effects on the movement of distant atoms within the protein due to interactions through hydrogen bonds (6). Despite their abundance and importance in protein structure, metal ions can be difficult to accurately parameterize for molecular dynamics (MD) simulations. Metal-bound proteins can contain complex coordination geometries with strong protein-metal interactions, charge transfer, polarization, and even varying coordination numbers (4). While some programs offer either bonded or non-bonded models to estimate the coordination geometry, many others cannot handle these ions and exclude them. While it may simplify the computational modeling of the given metalloprotein, excluding the metal ions may also affect the overall protein structure which could cause computational results to deviate from the experimental observations.

The structure of ACE2 has some complexities that can make it difficult to accurately model. In particular, ACE2 contains a Zn ion at the angiotensin catalytic site, which is approximately 22 Å away from the S1 binding site. Zn is well-known for being difficult to model accurately due to its unusually strong and complex interactions with select amino acids (7). While a few software packages allow parameterization and computation of the Zn ion, many other widely used packages are not designed to incorporate Zn, making it enticing to exclude this ion from the model. Due to both difficulties in modeling and its distance from the S1 binding site, many computational studies have chosen to exclude Zn from their models of ACE2 (8). However, studies have shown that Zn can have large effects as far away as 34 Å, so its exclusion may negatively impact the accuracy of these models (9).

In our previous work we have explored the implications of Zn-bound angiotensin peptides on ACE2-S1 binding (10). Additionally, we have discussed the time-based dynamics of immunologically relevant protein systems (11) (12). In this study, we evaluate the importance of Zn within a model structure of the ACE2-S1 complex by examining its impact on structural stability and binding free energies.

## Methods

### Overview

To examine the differences attributed to the presence of Zn, we obtained crystal structures for the ACE2-S1 complex, parameterized/excluded Zn ion, and performed MD simulations for both Zn-bound and Zn-free versions of the complex. Structural stability and binding free energy could then be computed and analyzed using the results of the MD simulation.

### Modeling ACE2-S1 Complexes

The experimentally-derived structure for the ACE2-S1 complex, chains A and E from 6M0J (13), was obtained from RCSB PDB (14). The Zn-free model structure was created by deleting Zn from one copy of the structure. Amber parameters, neutralizing ions (Na+ and Cl-), and a waterbox were generated for the Zn-free complex using the LEaP program (15).

The Zn-bound protein complex was created by parameterizing Zn in the other copy of the ACE2-S1 complex. To parameterize Zn with the appropriate coordination geometry, Amber force field parameters were generated for Zn and each coordinated residue using the bonded model in MCPB.py (1). In this method, force field parameters were generated with the Empirical method. During restrained electrostatic potential (RESP) charge fitting, the charges on the heavy backbone atoms of coordinated residues were constrained. Additionally, to convert from CHARMM to Amber notation and estimate residues’ protonation states at a pH of 7.0, Bio3D (16) and H++ (17) were used. Density functional theory was also employed, via GAMESS-US (13), to calculate the charges of atoms within the coordinating residues. Similar to the Zn-free complex, the LEaP program was used to add neutralizing salt and a waterbox with 10 Å before molecular dynamics simulations (15).

### Molecular Dynamics Simulations

After parameterizing both the Zn-bound and Zn-free complexes, MD simulations were performed in NAMD (18) to minimize energy and then to explore structural differences. Minimization occurred for 30,000 iterations using a step size of 1 fs at 310 K, controlled through Langevin dynamics with damping coefficient of 5 ps^-1^, and 1 atm, controlled with a Langevin piston with an oscillation period of 100 fs and damping timescale of 50 fs. Before moving to the next step, we checked to ensure that both total protein root mean square deviation (RMSD) and total energy had stabilized using the appropriate VMD plugins, RMSD and NAMD Energy (19). We then performed a constant volume MD simulation of each complex with a step size of 1 fs at 310 K, controlled through Langevin dynamics. A 2 ns constant volume equilibration MD simulation was followed by a 50 ns MD production run. For all MD simulations Particle mesh Ewald (PME) was used to account for electrostatic interactions.

### Analysis of Structural Stability and Binding Free Energy

Once the 50 ns production runs were complete, they could be used to analyze the structural stability and binding free energy of both the Zn-bound and Zn-free complexes. To evaluate structural stability, RMSD, root mean square fluctuations (RMSF), and secondary structure of each complex during the 50 ns MD simulation were visualized with VMD plugins. The binding free energy of each ACE2-S1 complex was calculated employing the MM/PBSA method, using the CaFE plugin for VMD (20). To calculate the solvent accessible surface area (SASA), APBS was used (21). Due to the system size and length of the MD simulation, binding free energy was computed with a step size of 0.45 ns. The solvent dielectric constant was kept at 80 while internal dielectric constants of 1, 2, 4, and 6 were used. As previously shown by Fatouros et al., larger internal dielectric constants are required for polar or charged binding sites (10).

## Results

We explored potential changes in the structural stability of the ACE2-S1 complex by computation of RMSD and root mean structure fluctuation (RMSF). In addition to the overall complex, measurements were made for the interacting residues (IR), defined as the residues within 3.5 Å of the opposite protein within the unmodified 6M0J crystal structure. The lower RMSD values of IRs compared with the overall protein reflects their higher inter- and intramolecular interactions, restricting their ability to move. From Figure 1A, it appears that the Zn-bound complex has slightly lower average RMSD values compared to the Zn-free complex, for both ligand and receptor IR and a greater average RMSD value for the overall complex. This is supported by Figure 1B-C, which shows that nearly all residues within the Zn-bound complex have slightly greater RMSF values than their counterparts within the Zn-free complex. As expected, the Zn-bound complex shows lower RMSD values than the Zn-free complex at residues with at least one atom within 15 Å of Zn (Figure S1). However, beyond this distance from Zn, the Zn-bound complex shows a greater RMSD than the Zn-free complex.

**Figure 1.**
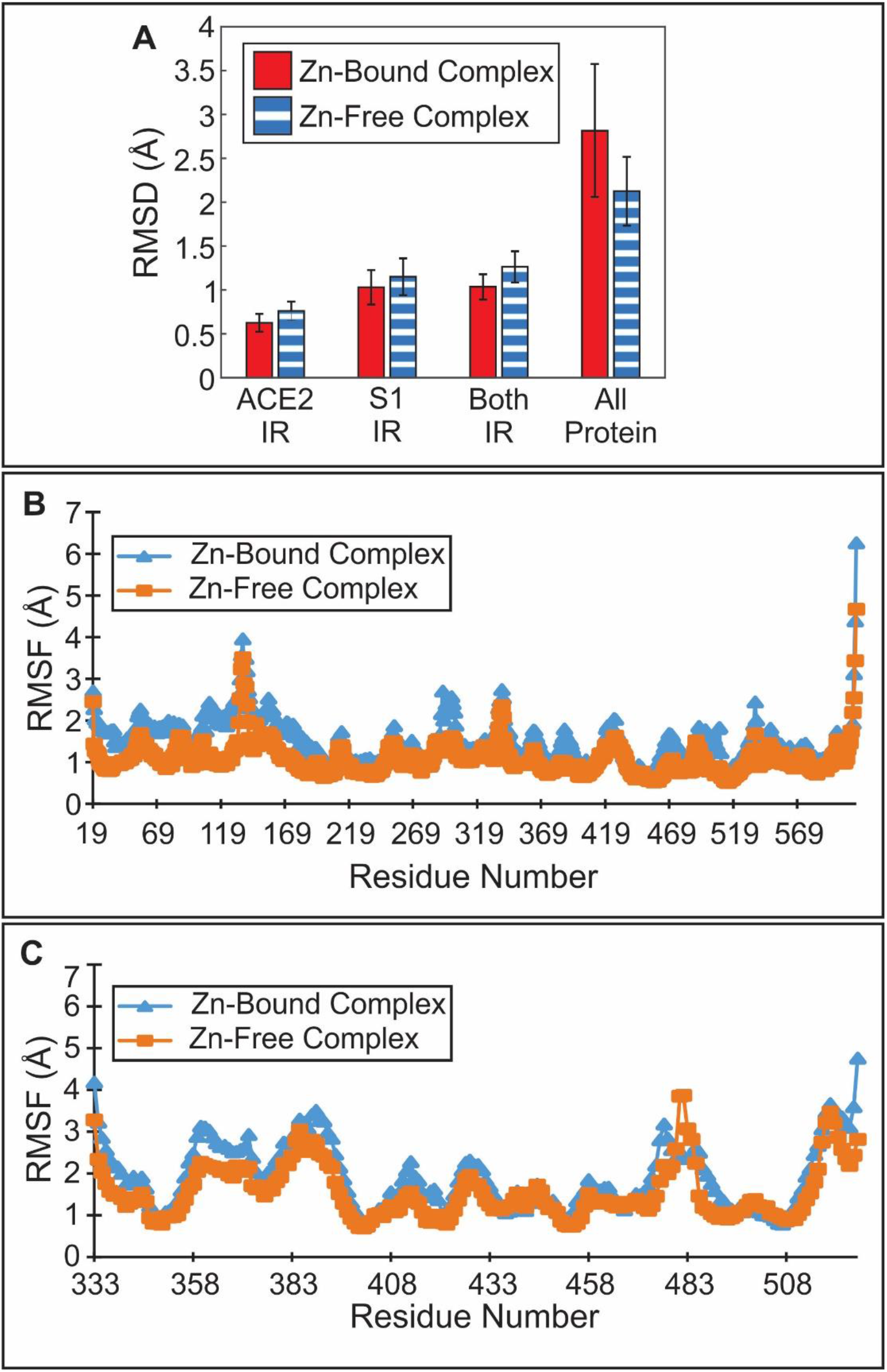
**A** Average RMSD values of all residues in the ACE2-S1 protein complex and interacting residues (IR) within ACE2, S1, or both. The values from the Zn-bound and Zn-free form of ACE2-S1 complex were computed for each condition. **B**-**C** RMSF for the residues within ACE2 (**B**) and S1 (**C**).

To investigate why the Zn-bound complex was more stable at the IR yet somewhat less stable overall, we examined the time-based secondary structures of each complex. A few differences are noticeable between the Zn-bound and Zn-free complexes (Figure S2). Unstable turns near ACE2 residue 107 in the Zn-bound complex are replaced by stable α-helices within the Zn-free complex. A similar change is also seen near ACE2 residue 448, where the Zn-bound complex contains unstable coils or 3-10 helix and the Zn-free complex contains stable α-helices. The secondary structures of the IR are of particular interest as it could offer an explanation of the deviations observed in Figure 1, and also due to potential impact on computed binding free energies. The secondary structures of IR within ACE2 show little variation between the complexes, although residue 355 of ACE2 is found to spend more time in stable β-sheet form in the Zn-bound complex (Figure 2). Interestingly, more differences are noticed within the S1 protein’s secondary structure. In particular, S1 protein’s residues 449 and 496 appear to change from stable β-sheets in the Zn-bound complex to unstable turns and coils in the Zn-free complex (Figure 2). In contrast, S1 residue 417 in the Zn-bound complex appears to fluctuate between stable α-helices and unstable turns and coils, while the residue remains as a stable α-helix in the Zn-free complex (Figure 2).

**Figure 2.**
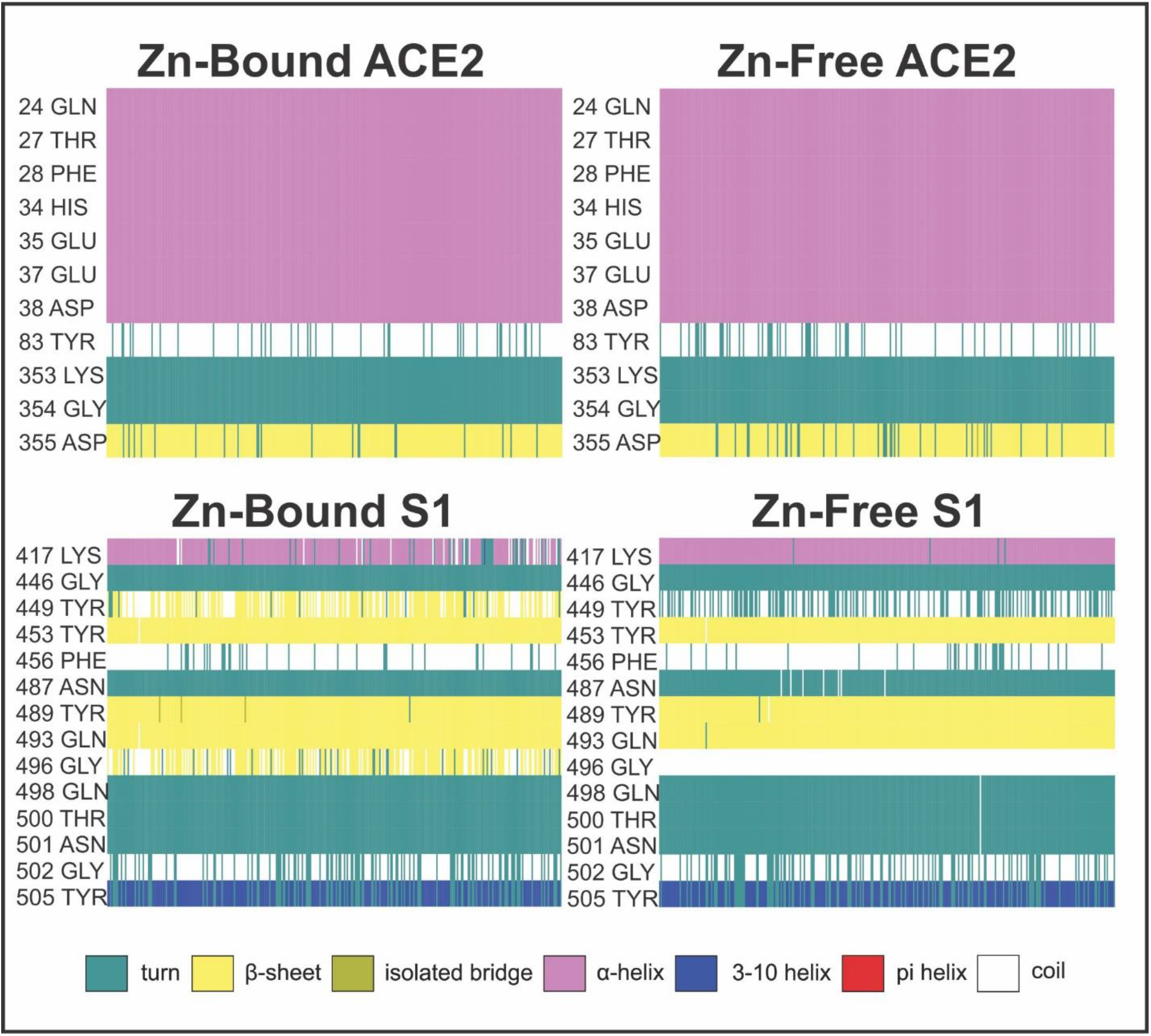
Time-based changes in the secondary structures of the interacting residues (IR) in ACE2 (top) and S1 (bottom), both in the Zn-bound (left) and Zn-free (right) complexes. The color at each horizontal position corresponds to a residue’s secondary structure at that time point in the MD simulation of 50 ns duration.

Since we observed differences in the stability of the IR and total protein between Zn-bound and Zn-free complexes, we wanted to investigate if this is translated into a difference in binding free energy. Binding free energies were computed for the ACE2-S1 complex both with and without Zn (Figure 3). A range of internal dielectric constants were considered during binding energy calculation in order to account for charged and polar IR, which require higher internal dielectric constants for accurate computation. We found that the binding free energies calculated with the internal dielectric constant of 1 align the most closely with the experimentally derived value of -11.83 kcal/mol (13). We also observed that the Zn-bound complex consistently has a stronger binding free energy compared to the Zn-free complex when higher internal dielectric constants were used in the computation.

**Figure 3.**
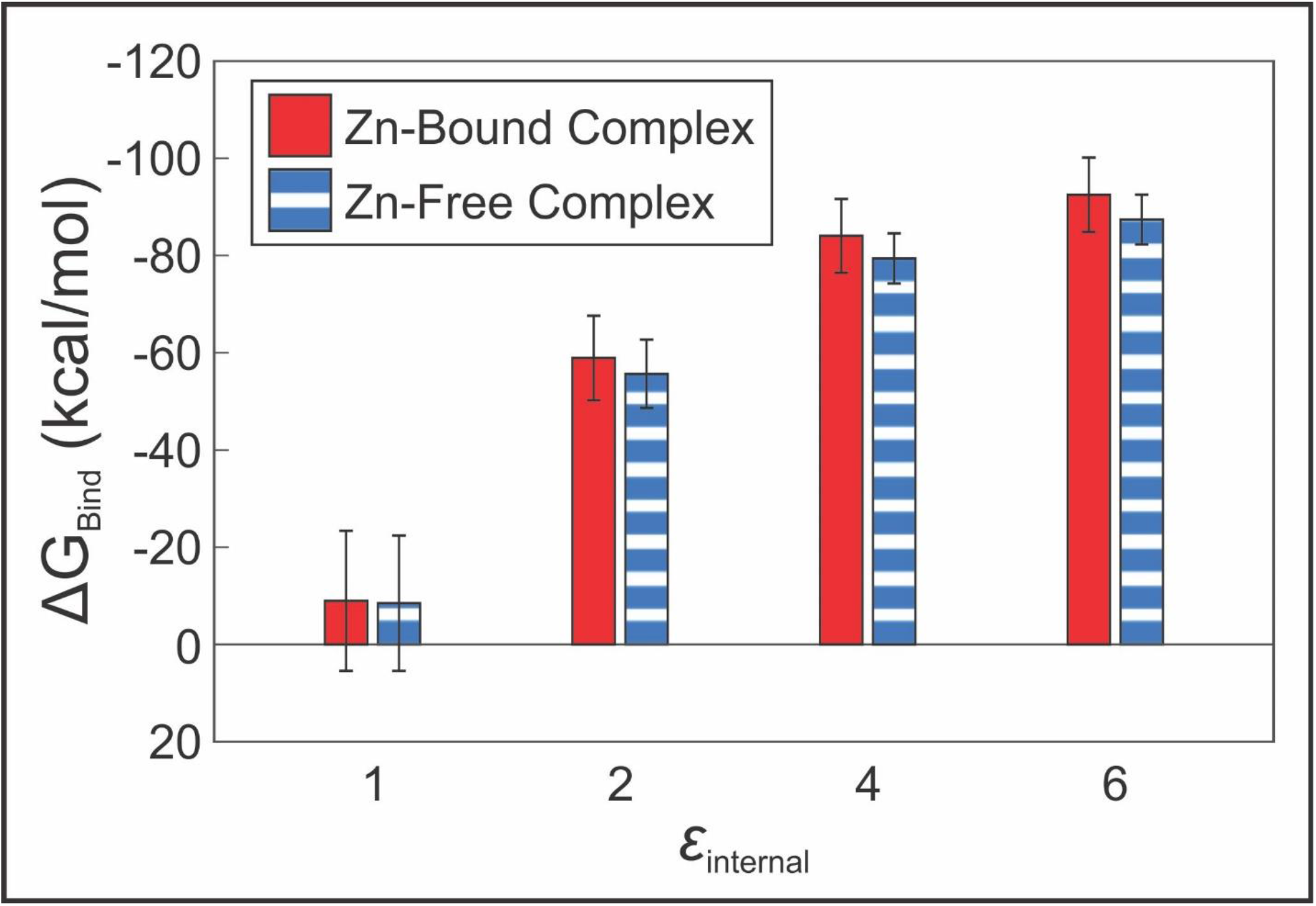
Binding free energies of the ACE2-S1 complexes computed using internal dielectric constants of 1, 2, 4, and 6. Both Zn-bound and Zn-free form of the complex were considered for computation.

## Discussion

Zn is expected to form strong connections to nearby residues within the protein and reduce movement, however, our results from MD simulation suggest an opposite effect of Zn on the overall structural stability of ACE2-S1 complex. We found that the protein within the Zn-bound complex has a greater average RMSD than the Zn-free complex, implying there is more movement occurring within the Zn-bound complex. However, there appear to be minor differences in the time-based secondary structures between these two complexes. As for example, ACE2 residues 107 and 448 changed from unstable turns and coils in the Zn-bound complex to more stable α-helices in the Zn-free complex (Figure S2).

Although the Zn-bound complex as a whole exhibits more movement, both the ACE2 and S1 IR appear more stable in the Zn-bound complex. In Figure 1A, both ACE2 and S1 IR show lower average RMSD in the Zn-bound complex, suggesting greater stability. While the secondary structures for the ACE2 IR demonstrate only minor differences, the secondary structures of the S1 IR are found to be more stable in the Zn-bound complex (Figure 2). For example, IR 449 and 496 in S1 change from more stable beta-sheets in the Zn-bound complex to unstable turns and coils in the Zn-free complex. While Zn is reported to have distant effects (9),at this time we do not fully understand why S1 residues located further away undergo changes even though much less changes are observed in the closer ACE2 residues. It is possible that the S1 residues are more susceptible to electrostatic and water-mediated hydrogen-bond interactions induced by the presence/absence of Zn, however, the reason for such susceptibility is yet to be understood. Another possible explanation is that when the Zn ion influences the movement of a pair of IR, any movement of the residue radially proximal to Zn needs to be compensated by a larger movement of a distal residue to maintain such interactions. This is supported by the finding that RMSD of residues within the Zn-bound complex increases with increase in the distance from the Zn ion (Figure S1). In contrast, in the Zn-free complex, little change in the RMSD value was observed for residues located within 10-20 Å away from Zn.

To investigate the apparently opposite effects on conformational stabilities of IR and overall protein, binding free energy was computed for the Zn-bound and Zn-free complexes. While computing the binding free energy, a variety of internal dielectric constants were used. While an internal dielectric constant of 1 is often the default value, it is not valid for all systems (22, 23). As the IR become more charged and polar, a greater internal dielectric constant is required to calculate accurate binding free energies (23). Thus, to account for differences in the polarity of the IR, a range of internal dielectric constants were used to calculate binding free energies. Based on the literature value for the binding free energy of ACE2-S1, -11.83 kcal/mol (13), using an internal dielectric constant of 1 yields the most accurate results. At this internal dielectric constant, the Zn-bound complex has a binding free energy that is nearly identical to the Zn-free complex. It is possible that, within the Zn-bound complex, the improved stability of IR could counteract the decreased stability of the other residues.

When using higher internal dielectric constants for computation of binding free energy, which are used for more polar protein-protein interfaces, the Zn-bound complex has a consistently stronger binding free energy. Some computational studies reported the calculated binding free energies of ACE2-S1 complex to be stronger than -30 kcal/mol (24), which would require an internal dielectric constant of greater than 1. It is possible that the S1 domain has been tuned through selective mutations to bind strongly to ACE2 with the involvement of Zn in this interaction, and thus excluding Zn creates a model that could underestimate the binding strength to S1. Since binding free energy is often used to evaluate potential inhibitors, differences due to Zn exclusion could potentially lead to inaccuracies in the assessment of their effectiveness.

## Conclusion

Our results show that the exclusion of Zn from the ACE2-S1 complex can influence its structural stability and may also have an impact on the calculated value of binding free energy. Although coordination to Zn is expected to stabilize nearby residues, we find its inclusion marginally reduces the overall stability of the protein complex with increased stability observed only in the IR. Interestingly, the inclusion of Zn was found to have a stronger influence on the secondary structure of S1 compared to the ACE2 receptor, even though S1 is located farther away from Zn. We have also observed that binding free energies for Zn-bound and Zn-free complexes are comparable at an internal dielectric constant of 1, however, the Zn-bound complex has stronger binding free energies at higher internal dielectric constants. Since binding free energy is often used to score the efficacy of inhibitors, relatively small differences in binding free energy values between the Zn-bound and Zn-free complex may lead to the inaccurate evaluation of candidate inhibitors. Our results could also provide some insights into the design of target inhibitors to other metalloprotein systems with the property of exhibiting differences in the binding free energy when the metal ions are excluded.

## Supplementary information

Supplementary Fig. S1 compares the RMSDs of all residues with at least one atom within the given radial distance from zinc in the zinc-bound complex. Supplementary Fig. S2 shows the time-based changes in secondary structures of all residues within the ACE2-S1 complex both with Zn and without Zn.

## Acknowledgements

The computations for this work were performed on the ACRES cluster. We would like to thank Clarkson University and the Office of Information Technology for providing computational resources and support that contributed to these research results. Additional computational resources for this grant were provided by the National Science Foundation under Grant No. 1925596. The authors acknowledge use of the following simulation and visualization software packages: 1) NAMD and 2) VMD: NAMD and VMD, developed by the Theoretical and Computational Biophysics Group in the Beckman Institute for Advanced Science and Technology at the University of Illinois, Urbana-Champaign.

